# geneslator: an R package for comprehensive gene identifier conversion and annotation

**DOI:** 10.64898/2026.03.30.714723

**Authors:** Giulia Cavallaro, Giovanni Micale, Grete Francesca Privitera, Alfredo Pulvirenti, Stefano Forte, Salvatore Alaimo

**Affiliations:** Istituto Oncologico del Mediterraneo (IOM), Via Penninazzo 7, 95029 Viagrande, Italy; Department of Clinical and Experimental Medicine, Bioinformatics Unit, University of Catania, Catania, Italy

**Keywords:** data integration, GeneID conversion, orthologs mapping, pathways mapping

## Abstract

**Motivation:** High-throughput sequencing generates large gene lists, making data interpretation challenging. Accurate gene annotation and reliable conversion between identifiers (e.g., gene symbols, Ensembl GeneIDs, Entrez GeneIDs) are essential for integrating datasets, conducting functional analyses, and enabling cross-species comparisons. Existing tools and databases facilitate annotation but often suffer from inconsistencies, missing mappings, and fragmented workflows, limiting reproducibility and interpretability.

**Results:** To address these limitations, we developed `geneslator`, an R package that unifies gene identifier conversion, orthologs mapping, and pathway annotation across eight model organisms (*Homo sapiens, Mus musculus, Rattus norvegicus, Drosophila melanogaster, Danio rerio, Saccharomyces cerevisiae, Caenorhabditis elegans, Arabidopsis thaliana*). `geneslator` provides an up-to-date, precise, and coherent framework that preserves data integrity, enables cross-species analyses, and facilitates robust interpretation of gene function and regulation, outperforming state-of-the-art gene annotation tools.

**Availability:** `geneslator` is available at https://github.com/knowmics-lab/geneslator.

## Introduction

High-throughput techniques have transformed biomedical research by enabling comprehensive sequencing of genomes, transcriptomes, and epigenomes. However, these approaches generate extensive gene lists, making data interpretation a significant challenge [1]. In addition, different databases, analysis tools, and computational pipelines often rely on distinct identifier systems, and inconsistencies between them can lead to data loss, misannotation, or incorrect biological interpretation. In this context, mapping among different gene identifier systems, such as gene symbols, Entrez Gene IDs, and Ensembl Gene Ids, is essential for transforming raw sequence data into meaningful biological knowledge, improving the understanding of genetic diseases, facilitating drug target discovery, and supporting the advancement of personalized medicine. Robust and up-to-date identifier conversion ensures interoperability across resources, enables the integration of heterogeneous datasets, and preserves the biological meaning of results throughout downstream analyses. As biological knowledge expands, mappings must be continuously updated to ensure compatibility with specialized analytical workflows. It is important to note that even modest increases in mapping efficiency can have a substantial impact on downstream analyses, as the successful conversion of a limited number of additional identifiers may significantly influence subsequent steps, including functional enrichment and orthologs mapping. Indeed, modern transcriptomics studies increasingly rely on comparative and functional analyses. Ortholog identification enables evolutionary comparisons and facilitates the translation of findings across model organisms, which is essential for validating discoveries in human disease contexts. Furthermore, pathway enrichment analysis - using frameworks such as Gene Ontology (GO) [2], Kyoto Encyclopedia of Gene and Genomes (KEGG) [3], Reactome[4] and Wikipathway [5] - contextualizes gene lists within biological processes, molecular functions and disease mechanisms, revealing coordinated regulatory patterns that individual gene annotations cannot capture. Despite their complementary nature, these analytical steps typically require multiple tools and databases, each with distinct formats, update schedules, and identifier systems, introducing inconsistencies and workflow fragmentation. Public bioinformatics resources, including NCBI (National Center for Biotechnology Information) [6], the Gene Nomenclature Committee (HGNC) [7], and the Ensembl Project [8], provide platforms for accessing gene information. While manual lookups are valuable for small-scale queries, they become impractical at the genome scale, where the process is time-consuming, error-prone, and must be repeated at each database release. Recently, several tools have been proposed to facilitate automated gene annotation, such as: (i) `biomaRt`, which provides programmatic access to BioMart databases for retrieving genomic annotations, (ii) organism-specific annotation packages (`org.*.*.db` packages), which offer curated, organism-level mappings between gene identifiers and functional annotations, (iii) `mygene` [9], a web-based API for fast gene annotation and identifier conversion, and (iv) `gprofiler2` [10], which enables functional enrichment analysis and identifier mapping through the g:Profiler platform. However, several challenges still persist, including discrepancies across resources and missing annotations that may compromise downstream analyses. Together, these limitations highlight the lack of a unified and up-to-date framework that simultaneously addresses identifier conversion, ortholog mapping, and functional annotation while preserving data integrity across downstream analyses. To overcome these limitations, we developed `geneslator`, a comprehensive R package that unifies gene identifier conversion, orthologs identification, and pathway mapping within a single, coherent framework. Our system is designed to maximize accuracy and precision while minimizing missing values. The enhanced precision of the tool stems from continuous incorporation of the latest gene annotations and mappings, ensuring a reliable, up-to-date conversion process. `geneslator` supports eight commonly studied organisms: *Homo sapiens, Mus musculus, Rattus norvegicus, Drosophila melanogaster, Danio rerio, Saccharomyces cerevisiae, Caenorhabditis elegans*, and *Arabidopsis thaliana*. It enables seamless cross-species comparisons and robust interpretation of gene function, regulation, and evolutionary conservation. By providing a unified analytical environment, it ensures methodological consistency and minimizes information loss across the entire workflow.

## Methods

Fig.1 illustrates the general workflow of `geneslator` R annotation package, which allows the user to query different types of data about a gene: annotations from general databases (symbol, aliases, full name, genetype), annotations from species-specific databases, functional annotations (pathways and gene ontologies), and orthologs.

**Fig. 1.**
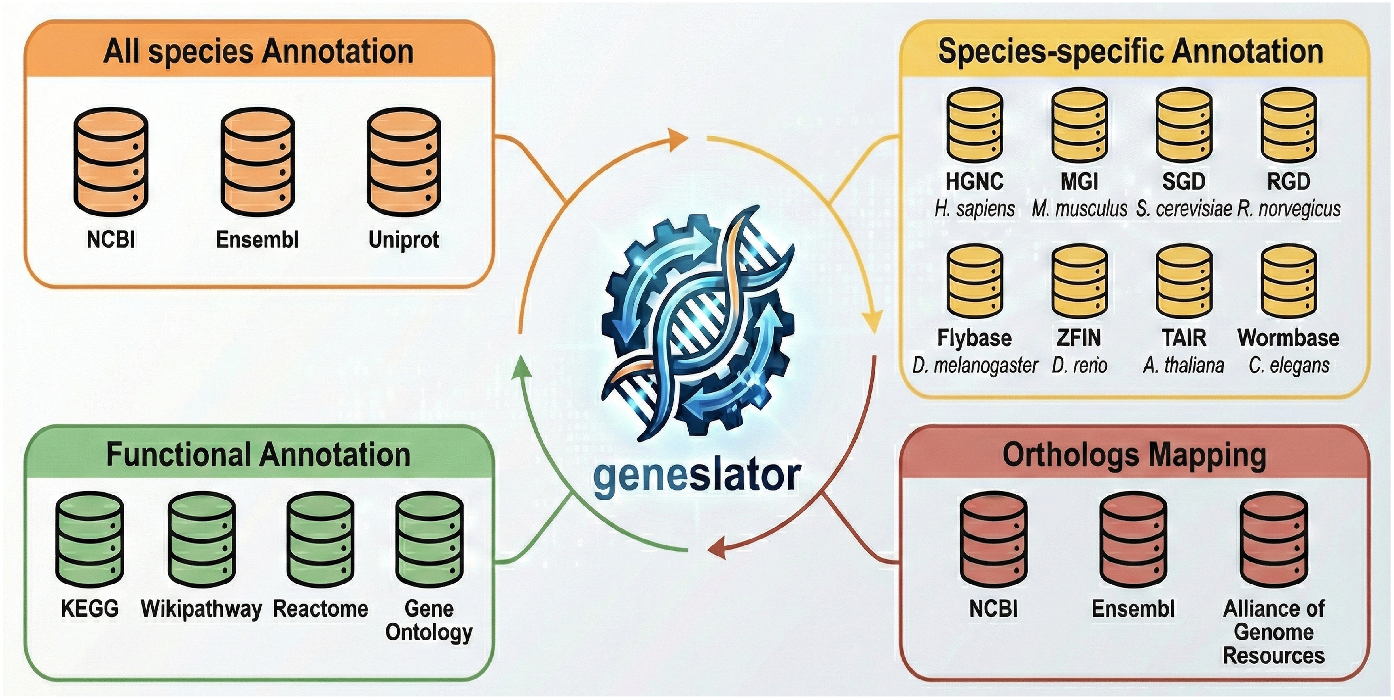
`geneslator` workflow. Gene identifiers are integrated from general and species-specific annotation sources, mapped to orthologs across model organisms, and linked to pathways via GO, KEGG, Reactome, and WikiPathways, providing a unified framework for functional and comparative analyses.

### Data preparation

As a preliminary step, annotation databases have been built for the following 8 model organisms: Human (*Homo sapiens*), Mouse (*Mus musculus*), Rat (*Rattus norvegicus*), Fly (*Drosophila melanogaster*), Zebrafish (*Danio rerio*), Yeast (*Saccharomyces cerevisiae*), Worm (*Caenorhabditis elegans*), and Arabidopsis (*Arabidopsis thaliana*). These databases integrate data collected from several sources, using the most up-to-date versions at the time of writing (up to December 15, 2025).

General information about a gene (symbol, aliases, full name, and genetype) was extracted from NCBI Gene [6] and Ensembl (v.115) [8]. Genetype represents the biotype classification of a gene (e.g., “protein-coding gene”, “non-coding RNA”, “pseudogene”, “lncRNA”). For *A*.*thaliana, C*.*elegans, D*.*melanogaster*, and *S*.*cerevisiae*, we have also extracted locus tag identifiers. Integration of NCBI and Ensembl data for the same gene was done by prioritizing NCBI information, as we noticed that Ensembl data were sometimes inaccurate across several organisms. For example, Ensembl associates id ENSG00000232654 to human gene FAM136BP, linking this information to entry HGNC:21110 in HGNC database [7]. However, in NCBI, HGNC:21110 is associated to NCBI Gene Id 387071, which correspond to the official gene symbol TIMCCP1, while FAM136BP is reported only as an alias of TIMCCP1. Similarly, id ENSDARG00000056314 is linked to zebrafish gene a2ml in Ensembl, while in NCBI Gene the same id is associated to the official gene symbol provided by Zebrafish Nomenclature Committee (ZNC) a2m2e, of which a2ml is an alias.

Identifiers of a gene include Entrez GeneIDs (taken from NCBI), Ensembl GeneIDs (taken from NCBI and Ensembl), Uniprot IDs of its proteins (taken from Uniprot (v.October 2025) [11]) and species-specific identifiers, coming from the most popular species-specific genome database, namely HGNC [7] for Human, MGI [12] for Mouse, RGD [13] for Rat, SGD [14] for Yeast, WormBase [15] for Worm, FlyBase [16] for Fly, ZFIN [17] for Zebrafish and TAIR [18] for Arabidopsis. Again, NCBI and Ensembl data referring to the same gene were integrated prioritizing the information coming from NCBI. In Zebrafish, we have also used HCOP [19] to link Ensembl GeneID to Gene symbols, because Ensembl’s annotations for this organism were often incorrect. In this case, we have given the highest priority to NCBI, followed by HCOP and Ensembl. In our annotation databases, we have also integrated old discontinued and replaced gene identifiers from NCBI gene and Ensembl (from v.100 to v.114). These archived identifiers have been stored in different columns with respect to current identifiers.

Genes’ orthologs have been retrieved from NCBI, Ensembl and AllianceGenome [20]. AllianceGenome provides a comprehensive orthology resource by consolidating predictions from 11 databases, namely OMA [21], SonicParanoid [22], InParanoid [23], Ensembl Compara [24], OrthoInspector [25], PhylomeDB [26], PANTHER [27], OrthoFinder [28], Hieranoid [29], ZFIN [17] and HGNC[7]. For Human, we also collected data from HCOP [19]. Orthologs have been represented by their Gene symbols and integration of orthologs data referring to the same gene has been done according to the following order: NCBI, HCOP (for Human), AllianceGenome and Ensembl.

Functional annotation data for a gene include pathways collected from KEGG Pathways [3], Reactome [4] and Wikipathways [30] and gene ontologies taken from GO [2]. For pathways, we extracted both their ids and their names, while for gene ontologies we extracted GO IDs, full names, types (biological process, cellular component or molecular function) and evidence codes of gene annotations.

Finally, annotation databases resulting from the integration of gene data have been built as SQLite objects using the `AnnotationForge` R package (v.1.48.0) [31].

### Package description

`geneslator` package is inspired by Bioconductor annotation R packages^1^. These packages contain organism-specific SQLite annotation databases that can be queried using a set of standard functions defined in the AnnotationDbi R package [32]. The most popular ones are `select` and `mapIds`, which link a list of gene identifiers (called keys) to gene’s associated data in other columns of the annotation database.

Although all these packages use the same functions to query annotation databases, each package is based on a single organism and databases are often centered on information coming from a single source, potentially missing other relevant data for gene annotation. For example, human annotation package `org.Hs.eg.db` relies only on data coming from NCBI Gene [6].

To overcome these limitations, `geneslator` generalizes the behavior of these annotation packages. Firstly, it provides a unique framework for querying annotation databases of multiple organisms, avoiding the usage of different R packages, one for each organism. Secondly, `geneslator` customizes query functions `select` and `mapIds` in order to take into account: i) the aliases of a gene when the user wants to map its symbol *X* to other data but *X* is not present in the annotation database as a symbol, ii) archive data when the user tries to link old identifiers of a gene to other data.

To ensure that `geneslator` operates with up-to-date annotation databases, these resources are updated monthly. At each update, the databases are bundled into a dedicated GitHub Release labeled using the `year.month` format (e.g., `2026.03`) and made available at https://github.com/knowmics-lab/geneslator/releases. Currently, the update process is performed manually, although we plan to implement an automated workflow for scheduled updates in the future.

To query an annotation database, the latter must be first imported by the user in the R global environment, using a specific function `GeneslatorDb`. When a database is imported for the first time, `GeneslatorDb` downloads it from the most recent GitHub Release and stores it in the local cache of the user’s installation folder of the package, so that each subsequent import of the database will only read the latter from the cache folder. To ensure that the user always works with the most updated version of the annotation databases present in the Github release, a versioning check system has been integrated in the package. More specifically, if the database is present but a newer version is available, the user will be asked to update it. If the answer is yes, the updated annotation database will be downloaded and stored in the local cache. To further support reproducibility, the function `GeneslatorDb` can be also used to import an older release of the annotation databases (e.g. release “2025.12”). This allows users to download and use specific previous versions of the database. This enables the exact replication of prior experiments when consistent annotations across analyzes are required.

`geneslator` has been submitted to Bioconductor and is currently under review. It is downloadable at https://github.com/knowmics-lab/geneslator and can be installed using R package `devtools`. Here, we show an usage example of `geneslator` for *H*.*sapiens*:

~~~
GeneslatorDb(“Homo sapiens”)
result <-geneslator::select(org.Hsapiens.db,
                              keys = df$symbol,
                              columns = “ENTREZID”,
                              keytype = “SYMBOL”)
~~~

### Benchmarking

In all 8 model organisms supported by `geneslator`, we performed several experiments to evaluate: i) the ability of `geneslator` to correctly map different types of data about genes, ii) how `geneslator`’s annotations influence other downstream tasks, such as functional annotation of genes. Table 1 lists all gene expression datasets used to perform the experiments, indicating, for each dataset, the relative organism, the year of publication, the type of identifier used for genes and the number of genes in the experiment.

**Table 1.** Datasets used for the experiments.

| Species | Dataset (Year) | ID Type (n) |
| --- | --- | --- |
| *H.sapiens* | GSE306533 [33] (2025) | Gene symbol (25427) |
|  | TCGA-BRCA [34] (2018) | Ensembl Gene ID (56602) |
|  | GSE179351 [35] (2021) | Entrez GeneID (26430) |
| *M.musculus* | GSE189541 [36] (2021) | Gene symbol (49492) |
|  | GSE272933 [37] (2024) | Ensembl Gene ID (56748) |
|  | GSE133054 [38] (2019) | Entrez GeneID (26301) |
| *R.norvegicus* | GSE237797[39] (2023) | Gene symbol (33014) |
|  | GSE143555[40] (2021) | Ensembl Gene ID (32662) |
|  | GSE134932[41] (2019) | Entrez GeneID (14004) |
| *C.elegans* | GSE49043[42] (2013) | Gene symbol (19065) |
|  | GSE184086[43] (2021) | Ensembl Gene ID (46904) |
|  | GSE292447[44] (2025) | Entrez GeneID (47632) |
| *D.melanogaster* | GSE64405[45] (2014) | Gene symbol (15340) |
|  | GSE145322[46] (2020) | Ensembl Gene ID (17697) |
|  | GSE64405[45] (2014) | Entrez GeneID (15241) |
| *D.rerio* | GSE306907[47] (2025) | Gene symbol (14969) |
|  | GSE274820[48] (2024) | Ensembl Gene ID (22350) |
|  | GSE121101[49] (2018) | Entrez GeneID (19640) |
| *S.cerevisiae* | GSE280426[50] (2024) | Gene symbol (6520) |
|  | GSE307461[51] (2025) | Ensembl Gene ID (6692) |
|  | GPL90[52] (2002) | Entrez GeneID (5805) |
| *A.thaliana* | GSE292401[53] (2025) | Gene symbol (26779) |
|  | GSE292906 [54] (2025) | Ensembl Gene ID (33562) |
|  | GPL198[52] (2002) | Entrez GeneID (22251) |

In a first set of experiments, we used `geneslator` to map different types of gene identifiers. Each experiment consists of mapping a list of input keys to output values, where keys and values can be Gene symbols, Ensembl GeneIDs or Entrez GeneIDs. This yields a total of six different types of tests that ensure comprehensive coverage of real-world use cases. To evaluate the quality of mapping, we measured the following parameters:

- *One-to-one mapped keys*: number of keys that are mapped with only one value. A higher proportion of uniquely mapped keys indicates a more accurate and reliable
- mapping, as it reflects the resolution of unambiguous gene identifier.
- *Unmapped keys*: number of keys that are unmapped. A lower proportion of unmapped keys indicates a more complete and robust mapping.
- *One-to-many mapped keys*: number of keys that are mapped to two or more values. A lower number of such keys generally reflects higher mapping specificity and reduced ambiguity, although multiple values associated with the same key may arise from genuine gene identifier ambiguities.
- *Mapping multiplicity*: mean number of values associated with keys that are mapped to two or more values. Like the number of one-to-many mapped keys, mapping multiplicity measures the specificity of the mapping, but is less influenced by the presence of few genes which genuinely have lots of associated identifiers.

We compared `geneslator` with the following R packages: `biomaRt` (v.2.67.0) [55], `mygene` (v.1.46.0) [9] and `gprofiler2` (v.0.2.4) [10]. We also used species-specific packages such as `org.Hs.eg.db` (v.3.22.0) [56], `org.Mm.eg.db` (v.3.22.0) [57], `org.Rn.eg.db` (v.3.22.0)[58], `org.Dm.eg.db` (v.3.22.0) [59], `org.Ce.eg.db` (v.3.22.0) [60], `org.Dr.eg.db` (v.3.22.0)[61], `org.Sc.sgd.db` (v.3.22.0) [62] and `org.At.tair.db` (v.3.22.0) [63]. To assess the performance of `geneslator` with respect to the other packages, we first built, for each experiment *E* and tool *T*, a boolean vector that indicates, for each key *k*, if *k* was mapped or not by *T* in *E*. Next, we calculated the *Jaccard similarity* and the *FDR of Fisher’s exact test* between the boolean vectors of `geneslator` and the other tools in each test.

In a second set of experiments, we tested the ability of `geneslator` to retrieve human orthologs in all model organisms supported by the package (except *A*.*thaliana*, as it is a plant species), starting from Gene symbols, Entrez IDs or Ensembl Gene IDs. In each test, we calculated the percentage of input genes that were successfully mapped to human orthologs. Higher mapping percentages indicate improved tool performance in capturing orthologs relationships across species. We compared `geneslator` with the following widely used orthologs mapping R packages: `orthogene` (v.1.16.1)[64], `homologene` (v.1.4.68.19.3.27)[65], `babelgene` (v.22.9)[66], and `gprofiler2` (which offers orthology mapping via the `g:Profiler` service)[67].

Finally, we evaluated the impact of using `geneslator` as an identifier mapping tool on downstream tasks, such as pathway annotation. Specifically, we annotated genes with KEGG pathways using the R package `clusterProfiler` [68] and its function `enrichKEGG`, which requires as input a set of gene identifiers of varying type depending on the organism used (e.g. Entrez GeneID for *H*.*sapiens*, Ensembl GeneID for *A*.*thaliana* and Gene locus for *C*.*elegans*). If needed, before performing pathway annotation, the user must run `clusterProfiler`’s `bitr` function to map its set of gene identifiers to the required type, starting from a specified annotation database. In our experiments, we first ran `bitr` function using either `org.*.*.db` or `geneslator` annotation databases, starting from all possible types of gene identifiers (excluding the type required by `enrichKEGG`). Next, we performed pathway annotation with `enrichKEGG` and, in each test and for each pathway *p*, we calculated the difference in the number of successfully mapped genes in *p* when starting from `geneslator` and `org.*.*.db` databases. A one-sample Wilcoxon signed-rank test was run to evaluate whether these differences were significantly greater than zero, using a one-sided alternative hypothesis (*α* = 0.05). To evaluate `geneslator` mapping performance in downstream analysis in real-life cases we also performed differential expression analysis on three datasets (human-TRACTISS clinical trial [69], mouse-GSE189541 [36] and zebrafish-GSE274820 [48]). Differential expressed genes (DEGs) have been calculated employing R packages limma [70] and edgeR [71]. Genes have been considered DEGs if *p* was lower than 0.05. All experiments and statistical analyses were performed in R (version 4.5.1). All R codes and datasets employed during testing can be found at https://github.com/knowmics-lab/geneslator-evaluation. Annotation databases used for the experiments were taken from release “2025.12” (https://github.com/knowmics-lab/geneslator-data/releases/tag/2025.12).

## Results

### Identifiers mapping

Table 2 reports the performances of `geneslator`, `biomaRt`, `org.Hs.eg.db`, `mygene` and `gprofiler2` for mapping gene identifiers (Gene symbols, Ensembl GeneIDs and Entrez GeneIDs) in *H*.*sapiens*.

**Table 2.** Performance results of geneslator and the compared tools for gene identifiers mapping in *Homo sapiens*.

| Mapping | *Homo sapiens* |  |  |  |  |  |
| --- | --- | --- | --- | --- | --- | --- |
|  |  | geneslator | biomaRt | org.Hs.eg.db | mygene | gprofiler2 |
| Gene symbol<br>↓<br>Entrez GeneID | One to one | **99.94%** | 78.6% | 98.05% | 97.86% | 79.68% |
|  | Unmapped | **0%** | 20.62% | 1.94% | 2.13% | 18.69% |
|  | One to many<br>(Multiplicity) | 0.06%<br>(2.87) | 0.78%<br>(7.63) | 0.01%<br>(2) | 0.01%<br>(2) | 1.63%<br>(4.42) |
|  | Jaccard | - | 0.78 | 0.98 | 0.98 | 0.80 |
|  | Fisher-test fdr | - | **<0.001** | 1 | 1 | **<0.001** |
| Gene symbol<br>↓<br>Ensembl GeneID | One to one | 91.64% | 87.64% | 91.23% | 90.17% | **94.72%** |
|  | Unmapped | **1.16%** | 5.54% | 2.93% | 2.71% | 3.62% |
|  | One to many<br>(Multiplicity) | 7.2%<br>(2.7) | 6.82%<br>(2.71) | 5.84%<br>(2.83) | 7.12%<br>(2.7) | 1.66%<br>(2.12) |
|  | Jaccard | - | 0.96 | 0.96 | 0.98 | 0.87 |
|  | Fisher-test fdr | - | 1 | **0.004** | 0.55 | **<0.001** |
| Entrez GeneID<br>↓<br>Ensembl GeneID | One to one | 91.16% | 87.67% | 90.92% | 90.56% | **92.56%** |
|  | Unmapped | **0.57%** | 5.13% | 1.86% | 2.6% | 5.18% |
|  | One to many<br>(Multiplicity) | 8.27%<br>(2.84) | 7.2%<br>(3) | 7.22%<br>(3) | 6.84%<br>(2.95) | 2.26%<br>(6.77) |
|  | Jaccard | - | 0.93 | 0.96 | 0.96 | 0.83 |
|  | Fisher-test fdr | - | **<0.001** | **0.004** | **0.02** | **<0.001** |
| Entrez GeneID<br>↓<br>Gene symbol | One to one | **100%** | 92.58% | 99.8% | 99.8% | 91.61% |
|  | Unmapped | **0%** | 6.93% | 0.2% | 0.2% | 7.83% |
|  | One to many<br>(Multiplicity) | 0%<br>(-) | 0.49%<br>(2.14) | 0%<br>(-) | 0%<br>(-) | 0.57%<br>(2.12) |
|  | Jaccard | - | 0.92 | 1 | 0.99 | 0.91 |
|  | Fisher-test fdr | - | **<0.001** | 1 | 1 | **<0.001** |
| Ensembl GeneID<br>↓<br>Gene symbol | One to one | **98.92%** | 73.38% | 61.70% | 77.03% | 70.78% |
|  | Unmapped | **0.05%** | 26.62% | 37.29% | 22.93% | 29.21% |
|  | One to many<br>(Multiplicity) | 1.03%<br>(2.04) | 0%<br>(-) | 1.01%<br>(6.61) | 0.04%<br>(2.17) | <0.01%<br>(2) |
|  | Jaccard | - | 0.72 | 0.63 | 0.76 | 0.72 |
|  | Fisher-test fdr | - | 1 | **<0.001** | **<0.001** | **<0.001** |
| Ensembl GeneID<br>↓<br>Entrez GeneID | One to one | **75.03%** | 44.02% | 61.71% | 62.42% | 43.99% |
|  | Unmapped | **24.28%** | 55.14% | 37.29% | 37.54% | 55.2% |
|  | One to many<br>(Multiplicity) | 0.69%<br>(2.06) | 0.84%<br>(7.47) | 1.01%<br>(6.61) | 0.04%<br>(2.17) | 0.81%<br>(7.67) |
|  | Jaccard | - | 0.58 | 0.82 | 0.82 | 0.58 |
|  | Fisher-test fdr | - | **<0.001** | **<0.001** | **<0.001** | **<0.001** |

In almost all experiments, `geneslator` consistently achieved the highest proportion of mapped genes and the lowest proportion of missing identifiers. In mapping Ensembl GeneID to Gene symbol, `geneslator` achieved a one-to-one mapping rate of 98.92%, substantially outperforming `biomaRt` (73.38%), `org.Hs.eg.db` (61.70%), `mygene` (77.03%) and `gprofiler2` (70.78%). Notably, the proportion of unmapped identifiers for `geneslator` was limited to 0.05%, whereas other tools exhibited markedly higher missing rates, ranging from 22.93% to 37.29%. Fisher’s exact test confirmed that the differences in mapping performance between `geneslator` and `org.Hs.eg.db`, `mygene`, and `gprofiler2` were statistically significant (FDR*<*0.001). A similar pattern was observed for Ensembl GeneID-to-Entrez GeneID mapping, where `geneslator` retained a one-to-one mapping rate of 75.03% with respect to 44.02% for `biomaRt` and 43.99% for `gprofiler2`. Some benchmark tools were consistently associated with substantially higher proportions of unmapped identifiers, exceeding 37% in several instances. All pairwise comparisons with competing tools showed statistically significant differences (FDR*<*0.001). For Entrez GeneID-to-Ensembl GeneID and Entrez GeneID-to-Gene symbol mappings, `geneslator` again demonstrated near-complete coverage, with one-to-one mapping rates of 91.16% and 100%, respectively, and negligible levels of missing identifiers. In particular, Entrez GeneID-to-Gene symbol mapping using `geneslator` resulted in no unmapped identifiers. The unmapped rate for Entrez GeneID-to-Ensembl GeneID mapping was only 0.57%, substantially lower than all competing tools (ranging from 1.86% to 5.18%). Statistical tests revealed significant differences between `geneslator` and `biomaRt` (FDR*<*0.001), `org.Hs.eg.db` (FDR=0.004), `mygene` (FDR=0.02) and `gprofiler2` (FDR*<*0.001) for Entrez GeneID-to-Ensembl GeneID mapping. Mappings from gene symbols to the other identifiers further confirmed the robustness of `geneslator`. Gene symbol-to-Ensembl Gene ID and Gene symbol-to-Entrez GeneID mappings achieved one-to-one mapping rates of 91.64% and 99.94%, respectively, with unmapped identifiers consistently below 1.16%. In contrast, alternative tools showed substantially higher missing rates, reaching up to 20.62% in Gene symbol-to-Entrez GeneID test and displaying higher levels of one-to-many mappings, suggesting an increased ambiguity. For Gene symbol-to-Ensembl GeneID test, statistically significant differences were observed when comparing `geneslator` to `org.Hs.eg.db` (FDR=0.004), and `gprofiler2` (FDR*<*0.001). Similarly, Gene symbol-to-Entrez GeneID mapping showed significant differences between `geneslator` and `biomaRt` (FDR*<*0.001) as well as `gprofiler2` (FDR*<*0.001).

In Table 3 we report performance results for identifiers mapping in *M*.*musculus*.

**Table 3.** Performance results of geneslator and the compared tools for gene identifiers mapping in *Mus musculus*.

| *Mus musculus* |  |  |  |  |  |  |
| --- | --- | --- | --- | --- | --- | --- |
| Mapping |  | geneslator | biomaRt | Org.Mm.eg.db | mygene | gprofiler2 |
| Gene symbol<br>↓<br>Entrez GeneID | One to one | **78.16%** | 47.77% | 70.68% | 70.65% | 47.81% |
|  | Unmapped | **21.77%** | 51.77% | 29.32% | 29.35% | 51.25% |
|  | One to many<br>(Multiplicity) | 0.07%<br>(2.14) | 0.46%<br>(3.14) | 0%<br>(-) | <0.01%<br>(2) | 0.94%<br>(4.79) |
|  | Jaccard | - | 0.61 | 0.91 | 0.91 | 0.62 |
|  | Fisher-test fdr | - | **<0.001** | 1 | 1 | **<0.001** |
| Gene symbol<br>↓<br>Ensembl GeneID | One to one | **95.66%** | 83.89% | 57.29% | 27.28% | 83.35% |
|  | Unmapped | **3.33%** | 15.65% | 42.4% | 72.69% | 15.31% |
|  | One to many<br>(Multiplicity) | 1.01%<br>(2.04) | 0.46%<br>(2.02) | 0.31%<br>(3.72) | 0.03%<br>(2.19) | 1.34%<br>(2.44) |
|  | Jaccard | - | 0.87 | 0.59 | 0.28 | 0.88 |
|  | Fisher-test fdr | - | 1 | **0.003** | 1 | **0.003** |
| Entrez GeneID<br>↓<br>Gene symbol | One to one | **99.99%** | 84.45% | 94.34% | 94.34% | 83.86% |
|  | Unmapped | **0.01%** | 15.35% | 5.66% | 5.66% | 15.95% |
|  | One to many<br>(Multiplicity) | 0%<br>(-) | 0.2%<br>(2.26) | 0%<br>(-) | 0%<br>(-) | 0.19%<br>(2.24) |
|  | Jaccard | - | 0.85 | 0.95 | 0.95 | 0.84 |
|  | Fisher-test fdr | - | **<0.001** | 1 | 1 | **<0.001** |
| Entrez GeneID<br>↓<br>Ensembl GeneID | One to one | **94.04%** | 84.48% | 88.02% | 39.5% | 84.44% |
|  | Unmapped | **4.88%** | 15.28% | 11.68% | 60.5% | 15.33% |
|  | One to many<br>(Multiplicity) | 1.08%<br>(2.02) | 0.24%<br>(2.25) | 0.3%<br>(2.2) | 0%<br>(-) | 0.23%<br>(2.27) |
|  | Jaccard | - | 0.89 | 0.93 | 0.41 | 0.89 |
|  | Fisher-test fdr | - | 0.14 | 0.41 | **0.02** | 0.14 |
| Ensembl GeneID<br>↓<br>Gene symbol | One to one | **99.3%** | 98.73% | 58.66% | 98.77% | 98.73% |
|  | Unmapped | **0.01%** | 1.27% | 40.77% | 1.23% | 1.26% |
|  | One to many<br>(Multiplicity) | 0.69%<br>(2.02) | 0%<br>(-) | 0.547%<br>(2.83) | 0.01%<br>(2) | 0.01%<br>(2) |
|  | Jaccard | - | 0.98 | 0.59 | 0.98 | 0.98 |
|  | Fisher-test fdr | - | 1 | **<0.001** | 1 | 0.82 |
| Ensembl GeneID<br>↓<br>Entrez GeneID | One to one | **75.21%** | 47.71% | 58.66% | 59.09% | 47.74% |
|  | Unmapped | **24.34%** | 51.82% | 40.77% | 40.91% | 51.9% |
|  | One to many<br>(Multiplicity) | 0.45%<br>(2.02) | 0.47%<br>(3.01) | 0.57%<br>(2.83) | 0.01%<br>(2) | 0.36%<br>(2.49) |
|  | Jaccard | - | 0.64 | 0.78 | 0.78 | 0.64 |
|  | Fisher-test fdr | - | **<0.001** | **<0.001** | **0.002** | **<0.001** |

`geneslator` consistently achieved the highest proportion of one-to-one mappings, ranging from 75.21% to 99.99%, depending on the identifier type. Notably, `geneslator` reached more than 99% of one-to-one mapped genes when mapping from Entrez GeneID to Gene symbol (99.99%) and from Ensembl GeneID to Gene symbol (99.3%), with no unmapped genes (0.01%). In contrast, other tools exhibited substantially lower one-to-one rates and higher proportions of unmapped identifiers. `geneslator` also demonstrated a marked reduction in unmapped genes compared to other tools. For example, in Gene symbol-to-Ensembl GeneID task, `geneslator` left only 3.33% of genes unmapped, whereas `biomaRt` and `gprofiler2` failed to map 15.65% and 15.31% of identifiers respectively. `org.Mm.eg.db` and `mygene` showed even higher unmapped rates of 42.4% and 72.69%. In Gene symbol-to-Entrez GeneID mapping, `geneslator` maintained a rate of unmapped identifiers of 21.77%, comparable to `org.Mm.eg.db` and `mygene` (29.32% and 29.35%), but substantially lower than `biomaRt` (51.77%) and `gprofiler2` (51.25%). Similar trends were observed in Ensembl GeneID-to-Entrez GeneID task, where `geneslator` achieved 75.21% one-to-one mappings with 24.34% unmapped identifiers, compared to `biomaRt` and `gprofiler2` which both exceeded 51% unmapped rates. Statistical comparison using Fisher’s exact test confirmed that the differences in mapping performance between `geneslator` and the other tools were highly significant in nearly all scenarios. For Gene symbol-to-Entrez GeneID mapping, significant differences were observed when comparing `geneslator` to `biomaRt` and `gprofiler2` (*FDR <* 0.001), as well as to `org.Mm.eg.db` and `gprofiler2` in the Gene symbol-to-Ensembl GeneID case (*FDR* = 0.003). Similarly, Entrez GeneID-to-Gene symbol mapping showed significant differences between `geneslator` and `biomaRt` (*FDR <* 0.001) as well as `gprofiler2` (*FDR <* 0.001). For Entrez GeneID-to-Ensembl GeneID mapping, significant differences were observed with `mygene` (*FDR* = 0.02). In Ensembl GeneID-to-Gene symbol test, `geneslator` significantly outperformed `org.Mm.eg.db` (*FDR <* 0.001). Finally, Ensembl GeneID-to-Entrez GeneID mapping showed significant differences between `geneslator` and all competing tools, with `biomaRt`, `org.Mm.eg.db` and `gprofiler2` (*FDR <* 0.001), and `mygene` (*FDR* = 0.002).

In Supplementary Tables S1-S6, we show performance results for identifiers mapping in the other six model organisms supported by `geneslator` (*D*.*rerio, S*.*cerevisiae, A*.*thaliana, R*.*norvegicus, D*.*melanogaster*, and *C*.*elegans*). `geneslator` consistently shows superior performance, with one-to-one mapping rates above 95% in most experiments, reaching near-perfect or perfect mapping (99%-100%) in Entrez GeneID-to-Gene Symbol tests for several species. The percentage of unmapped identifiers in `geneslator` was often below 1%, whereas in the other tools this percentage frequently exceeded 10%-20%. Statistical comparisons using Fisher’s exact test confirmed significant differences (*FDR <* 0.001) between `geneslator` and the other tools in many scenarios across all species, validating the robustness and superior coverage of `geneslator`’s gene identifier mapping strategy. Moreover, in certain species, some compared tools did not work properly or were unable to produce results. For example, in *S*.*cerevisiae*, `org.Sc.sgd.db` exhibits poor performance when mapping from Gene symbol, because the SYMBOL column is missing and there is a GENENAME column that does not correspond to the gene symbol (Table S2). In *A*.*thaliana*, no results were reported for `biomaRt`, as the latter does not support plant species (Table S3). In some mapping tests of *S*.*cerevisiae* and *A*.*thaliana* (Tables S2 and S3), we were unable to run `mygene` due to documented bugs in its code.

Overall, experiments on identifier mapping show that `geneslator` outperforms existing tools with a higher number of one-to-one mappings and a lower number of unmapped identifiers. In many scenarios, `geneslator` identified a small percentage of one-to-many mappings, sometimes higher than the compared tools. However, these cases were manually reviewed and cross-referenced against the relevant reference databases, confirming that they corresponded to valid multiple mappings. This suggests that some one-to-one mappings or unmapped identifiers in other tools actually represent missing mappings. Besides performance results, experiments on identifiers mapping also reveal some limitations of the compared tools that negatively affect their usability and interpretation of the results. For example, `biomaRt` always requires an active internet connection, which prevents its use in offline environments and limits reproducibility in restricted computing settings. Additionally, it does not provide annotations for certain widely studied organisms, such as *A*.*thaliana*, reducing its applicability in plant transcriptomics studies. `mygene` becomes considerably slow when mapping large sets of gene identifiers. Moreover, when a gene identifier cannot be mapped, `mygene` maps the input identifier to itself anyway. This behavior can lead to ambiguous or misleading results and requires additional post-processing steps to correctly identify unmapped genes. Like `mygene`, `gprofiler2` shows inconsistent behavior with identifiers that cannot be mapped. In several cases, genes are incorrectly mapped to multiple identifiers or in an inconsistent manner, leading to a higher number of one-to-many mappings and a lower number of unmapped genes. These inconsistencies introduce biases that may affect downstream analyses, such as functional annotations with pathways and gene ontologies. Altogether, these findings suggest that existing tools often follow different mapping strategies and error-handling mechanisms, reinforcing the need for robust, transparent and well-validated gene ID mapping approaches.

### Orthologs mapping

Table 4 shows the results achieved by `geneslator` and the compared tools in retrieving human orthologs in all model organisms supported by `geneslator`.

**Table 4.** Performance results of geneslator and the compared tools for orthologs mapping.

|  |  | geneslator | orthogene<br>(gprofiler) | orthogene<br>(babelgene) | homologene | gprofiler2 | babelgene |
| --- | --- | --- | --- | --- | --- | --- | --- |
| *M.musculus* | Gene symbol | **43.28%** | 36.93% | 36.99% | 33.47% | 36.93% | 35.44% |
|  | Ensembl GeneID | **38.49%** | 35.77% | \ | \ | 35.77% | 33.66% |
|  | Entrez GeneID | **77.66%** | \ | \ | 65.02% | 70.15% | 71.74% |
| *C.elegans* | Gene symbol | **43.21%** | 0.01% | 27.99% | 15.09% | 33.35% | 22.80% |
|  | Ensembl GeneID | **18.29%** | \ | \ | \ | 14.37% | 13.62% |
|  | Entrez GeneID | **17.99%** | \ | \ | 7.92% | 14.00% | 13.40% |
| *D.melanogaster* | Gene symbol | **58.26%** | \ | 44.71% | 29.13% | 42.54% | 38.64% |
|  | Ensembl GeneID | **49.60%** | \ | \ | \ | 39.14% | 41.58% |
|  | Entrez GeneID | **57.63%** | \ | \ | 29.34% | 45.76% | 48.37% |
| *R.norvegicus* | Gene symbol | **63.57%** | 51.89% | 53.10% | 46.00% | 51.89% | 51.11% |
|  | Ensembl GeneID | **63.73%** | 53.32% | \ | \ | 53.32% | \ |
|  | Entrez GeneID | **94.67%** | \ | \ | 87.43% | 85.15% | 91.59% |
| *D.rerio* | Gene symbol | **85.19%** | 72.35% | 83.17% | 55.59% | 72.35% | 80.28% |
|  | Ensembl GeneID | **80.30%** | 69.70% | \ | \ | 69.70% | 73.83% |
|  | Entrez GeneID | **84%** | \ | \ | 60.77% | 69.94% | 77.49% |
| *S.cerevisiae* | Gene symbol | **53.97%** | 40.49% | \ | 0.81% | 40.49% | 37.44% |
|  | Ensembl GeneID | **53.09%** | 40% | \ | 0.79% | 40% | 37.66% |
|  | Entrez GeneID | **60.95%** | \ | \ | 23.24% | 45.79% | 43% |

In all experiments `geneslator` consistently achieved the highest percentage of genes mapped with human orthologs, regardless of the starting species or identifier type (Gene symbol, Entrez GeneID or Ensembl Gene ID), with the highest rate of identifiers mapped with orthologs (94.67%) reached in *R*.*norvegicus*, starting from Entrez GeneID. In *C*.*elegans* and *D*.*melanogaster*, `geneslator` substantially outperformed `orthogene` and `gprofiler2`, which in some cases failed to retrieve meaningful orthologs mappings from Gene symbols. Similarly, in *D*.*rerio* and *S*.*cerevisiae* `geneslator` showed robust and balanced performance, providing higher and more consistent orthologs coverage compared to other tools, even in species where annotation incompleteness poses significant challenges. Overall, these results highlight `geneslator`’s ability to deliver reliable and comprehensive orthologs mapping across diverse species and identifier systems, making it a valuable resource for cross-species and comparative genomics analyses.

### Pathway mapping

Table 5 shows the performance of functional annotation with `clusterProfiler`, using either `geneslator` or `org.*.*.db` annotation databases for preliminary identifier mappings.

**Table 5.** Performance results for functional annotation with clusterProfiler, using either geneslator or org.*.*.db annotation databases for preliminar identifier mappings.

|  | ID type | Top-mapped pathways | p | Average number of extra mapped genes |
| --- | --- | --- | --- | --- |
| *H.sapiens* | Gene symbol → Entrez GeneID | 17.05% | <0.001 | 4.13 |
|  | Ensembl GeneID → Entrez GeneID | 9.94% | 0.09 | 1.77 |
| *M.musculus* | Gene symbol → Entrez GeneID | 69.25% | <0.001 | 4.9 |
|  | Ensembl GeneID → Entrez GeneID | 25.29% | <0.001 | 2.9 |
| *D.rerio* | Gene symbol → Entrez GeneID | 57.14% | <0.001 | 2.3 |
|  | Ensembl GeneID → Entrez GeneID | 49.71% | <0.001 | 2.54 |
| *S.cerevisiae* | Entrez GeneID → Ensembl GeneID | 9.26% | 0.002 | 1.2 |
| *A.thaliana* | Gene symbol → Ensembl GeneID | 94.89% | <0.001 | 20.79 |
|  | Entrez GeneID → Ensembl GeneID | 5.11% | 0.009 | 1.28 |
| *R.norvegicus* | Gene symbol → Entrez GeneID | 83.91% | <0.001 | 5.90 |
|  | Ensembl GeneID → Entrez GeneID | 93.97% | <0.001 | 8.65 |

In several organisms, `geneslator`-based mappings resulted in a significant higher number of genes successfully mapped to KEGG pathways compared to `org.*.*.db`. The rate of improvement was species- and identifier-dependent. In *H*.*sapiens*, mapping from Gene symbols to Entrez GeneID using `geneslator` increased gene coverage in 17.09% of pathways (*p <* 0.001), with an average gain of 4.13 genes per pathway, whereas no significant difference was observed when Ensembl GeneID identifiers were used (*p* = 0.09).

In Supplementary File 1, KEGG functional annotation for *H*.*sapiens* using gene symbols reveals marked differences in pathway ranking between the two tools. Notably, *Salmonella infection*, the top-ranked pathway according to `org.Hs.eg.db`, is ranked second by `geneslator`, while *Pathways of neurodegeneration – multiple diseases*, ranked first by `geneslator`, is placed third by `org.Hs.eg.db`. Similar discrepancies are evident for additional pathways. For instance, *Oxidative phosphorylation* ranks 34th with `geneslator` (GeneRatio: 130/7750, *p* = 1.54e-04), but declines to 53rd with `org.Hs.eg.db` (GeneRatio: 115/7700, *p* = 0.53), resulting in a shift from significant to non-significant enrichment. These differences are likely explained by the distinct numbers of genes assigned to pathways by each tool, which in turn influence pathway ranking and statistical significance in gene-ratio-based analyses.

In *M*.*musculus* and *D*.*rerio*, `geneslator` consistently improved pathway gene coverage for both Gene symbol and Ensembl GeneID inputs, with a relevant percentage of pathways, respectively 69.25% and 25.29%, showing increased gene mapping and highly significant enrichment results (*p <* 0.001 in all cases).

As reported in Supplementary File 2, KEGG functional annotation of *M*.*musculus* using gene symbols highlights marked differences between `geneslator` and `org.Mm.eg.db`, with 88 pathways showing opposite *p* behavior across the two analyses. Ranking the results by *p* further underscores these discrepancies: the pathway placed first by `org.Mm.eg.db` is ranked only 36th by `geneslator`. This difference is attributable to the larger number of genes mapped by `geneslator` to the PI3K-Akt signaling pathway, as well as to the higher total number of genes mapped to pathways in the analysis overall (`geneslator`: 10,539; `org.Mm.eg.db`: 9,003). Slight but statistically significant improvements were observed in *S*.*cerevisiae* and *A*.*thaliana* starting from Entrez GeneID, where `geneslator` increased the number of mapped genes in a smaller proportion of pathways. Interestingly, in *A*.*thaliana*, `geneslator` mapped 94.89% more pathways than `org.At.tair.db` with, on average, 20.79 extra genes mapped for each pathway starting from Gene symbol. The most relevant improvements in functional annotation were detected in *R*.*norvegicus*, where `geneslator` improved gene mapping in the majority of pathways for both Gene symbol (83.91%) and Ensembl GeneID (93.97%) identifiers, yielding a large average number of extra annotated genes per pathway. For *D*.*melanogaster* and *C*.*elegans*, we were not able to compare `geneslator` with `org.*.*.db`, because for these organisms `clusterProfiler`’s `enrichKEGG` function requires as input locus tag identifiers that are not provided by `org.*.*.db` packages. Moreover, for *S*.*cerevisiae*, we only report results for Entrez GeneID-to-Ensembl GeneID mapping, because in `org.Sc.sgd.db` the SYMBOL column is missing and the GENENAME column does not correspond to the gene symbol.

We further evaluated whether improvements in gene identifier mapping translate into tangible benefits for downstream analyses. We applied `geneslator` to the differentially expressed genes from the TRACTISS clinical trial [69] which investigates anti-B cell therapy in primary Sjögren syndrome. We used RNA-Seq data are publicly available (https://tractiss.hpc.qmul.ac.uk/) and performed pathway enrichment analysis. The results show that `geneslator` and org.Hs.eg.db are biologically concordant in this context, with 72/72 pathways shared and identical top-19 enriched terms. Nevertheless, `geneslator` exhibits markedly higher sensitivity for pathways of direct relevance to the TRACTISS biological setting: Tight junction (≃ 57 − *fold* lower *p*), RNA polymerase (≃ 186 − *fold*), RNA degradation (≃ 75 − *fold*), Motor proteins (≃ 340 − *fold*) and O-glycan biosynthesis (≃ 30 − *fold*). For these substantial statistical advantages were observed. These were driven by the recovery of 2-4 additional annotated genes per pathway. The RNA polymerase pathway deserves particular attention, as it displayed the largest shift in enrichment significance across all datasets examined (Δ = −18). This finding is biologically well supported, since rituximab, the therapeutic agent evaluated in the TRACTISS trial, induces extensive transcriptional reprogramming in salivary gland tissue. Accordingly, the ability of `geneslator` to identify this pathway with full statistical significance (*p* = 1.56 *×* 10^−4^ versus *p* = 2.90 *×* 10^−2^ with `org.Hs.eg.db`) has direct clinical relevance for the mechanistic interpretation of treatment response. Further details are reported in Supplementary File 3.

The same approach was applied to the identification of DEGs in GSE189541, comparing NSCHRas-shp53 mice with NPCTKO mice. This analysis identified 93 pathways shared by both methods. Overall, the two approaches showed the strongest agreement for the most biologically relevant pathways. The largest ranking shifts, observed for *ATP-dependent chromatin remodeling* (Δ = −12) and *alanine metabolism* (Δ = −8), involved pathways for which `geneslator` identified additional genes relative to `org.Mm.eg.db`, resulting in lower *p*. However, these pathways remained within the mid-to-high ranking range (positions 27– 56), where enrichment was either close to the significance threshold or not statistically significant. Further details are provided in Supplementary File 4.

For zebrafish, analysis of GSE274820 using DEGs identified by comparing IWR1-treated samples with DMSO controls yielded the same 20 pathways with both tools. At first glance, the main difference appears to be that `org.Dr.eg.db` identifies *ATP-dependent chromatin remodeling* as a significant pathway (*p* = 4.50 *×* 10^−3^), whereas `geneslator` does not. However, closer examination of the gene-level differences between the two analyses indicates that `org.Dr.eg.db` introduces mapping errors. Specifically, it incorrectly associates ENSDARG00000109926 and ENSDARG00000111536 with Entrez Gene ID 100148591, whereas `geneslator` correctly maps them to 570115 and 560527, respectively. Another error concerns ENSDARG00000099123, which is incorrectly assigned to 103908687 instead of 100006331. A further discrepancy involves ENSDARG00000100906, which is wrongly mapped to 100001004 rather than to the correct identifier, 100004510. Additional details are reported in Supplementary File 5.

Thus, even when biological conclusions are concordant, the more accurate and less ambiguous annotations provided by `geneslator` yield a more robust foundation for enrichment analysis, reducing mapping noise and allowing statistical significance to better reflect the underlying biology.

## Discussion

Experiments show that `geneslator` is able to uniquely map a higher number of gene identifiers than the existing annotation tools with a very low rate of unmapped identifiers. The superior accuracy of `geneslator` is due to the incorporation of NCBI and Ensembl archived identifiers into our annotation databases and the usage of gene aliases in searches involving gene symbols, which allows retrieval of previous names or synonyms associated with gene symbols. These features enhance the robustness of our approach by accounting for nomenclature changes over time, minimizing discrepancies in genes annotation across different datasets. Experimental results were consistent across all organisms and identifier types, and these differences had tangible effects on downstream analyses. As shown in our experiments, even small improvements in gene identifier conversion rates can translate int substantial changes in downstream functional analyses, because minor increases in annotation completeness may propagate and amplify across enrichment and interpretation steps, ultimately altering biological conclusions. For example, in transcriptomic workflows, incomplete or ambiguous gene mappings can lead to the exclusion of a substantial proportion of features, thereby influencing differential expression results, pathway enrichment analyses, and subsequent biological interpretation. Moreover, we identified cases in which `org…db` packages produced incorrect mappings (see Supplementary File 6), with the potential to mislead functional interpretation. By minimizing information loss at the annotation stage, `geneslator` preserves data complexity and improves the robustness and reproducibility of downstream analyses. Moreover, `geneslator` effectively overcomes issues related to outdated annotations, inconsistent databases and ambiguous Gene symbol usage, which remain major sources of error in bulk RNA-seq analysis pipelines. The dynamic update mechanism and rigorous validation framework implemented in `geneslator` ensure high annotation accuracy and long-term reliability. Importantly, `geneslator` is based on a unified, organism-agnostic framework that provides a standardized annotation pipeline and ensures consistent implementation across all supported species through a shared underlying architecture. Although the primary goal of our package is to improve gene identifiers mapping, we deliberately included in our annotation databases also functional data coming from databases such as Reactome, KEGG, GO and WikiPathways to enable users to map genes of interest and explore their associated biological functions. This feature is intended to facilitate functional analyses also for species and identifier types that are not supported by existing annotation resources. It is worth noting that integrating functional data into our annotation databases does not mean that `geneslator` is able to perform enrichment statistical analyses, as done by tools like `clusterProfiler`. Nevertheless, the output of functional annotation produced by `geneslator` is fully compatible with established enrichment workflows and can be readily integrated with these tools, allowing users to perform downstream enrichment analyses without loss of functionality. Overall, `geneslator` helps researchers to seamlessly integrate refined and up-to-date gene annotations into their analyses, regardless of the starting identifier system. By enhancing gene ID mapping through continuous updates, `geneslator` improves annotation coverage and quality, streamlines a critical step of bulk RNA-seq analysis, and ultimately enables more comprehensive and biologically meaningful interpretations of transcriptomic data, fostering a deeper understanding of genomic landscapes and complex biological processes.

## Conclusion

In this paper, we introduced `geneslator`, a novel and user-friendly R annotation package designed to address critical limitations in current annotation tools. `geneslator` integrates different types of gene-related data from several sources into a unique framework and empowers standard functions used to query annotation databases, by extending, if needed, the search to archived identifiers and gene aliases. As shown experimentally, `geneslator` is able to map a substantially larger number of gene identifiers than the state-of-the-art annotation tools, across several commonly used model organisms. This, in turn, improves the robustness and reproducibility of downstream tasks, such as differential expression analysis or pathway enrichment. `geneslator` package is under continuous development and in the future we aim at increasing the number of supported species, including less commonly studied organisms, with the goal of providing a more comprehensive and inclusive framework for gene identifier translation.

## Data Availability

Identifiers data that support the findings of this study have been deposited in https://github.com/knowmics-lab/geneslator-data.

## Author contributions statement

GC, GFP, GM, SA, and AP conceived the work. GC, GFP and GM conducted the experiments. GC, GFP, GM, SA AP, and SF analyzed the results. GM developed the R package. GC, GFP, GM and AP wrote the first draft of the manuscript. All the authors reviewed and approved the manuscript.

## Acknowledgments

The findings reported in this paper are derived from data produced by the TCGA Research Network, accessible at https://www.cancer.gov/tcga. This study was funded by the 2024/2026 Research Plan of University of Catania Pia.ce.ri (IMAGINE project).

---

**Table S1.** Performance results of geneslator and the compared tools for gene identifiers mapping in *D*.*rerio*.

|  |  | *Danio rerio* |  |  |  |  |
| --- | --- | --- | --- | --- | --- | --- |
| Mapping |  | geneslator | biomaRt | Org.Dr.eg.db | mygene | gprofiler2 |
| Gene symbol<br>↓<br>Entrez GeneID | One to one | **99.84%** | 90.57% | 94.11% | 93.58% | 72.9% |
|  | Unmapped | **0.01%** | 8.69% | 5.89% | 6.41% | 6.61% |
|  | One to many<br>(Multiplicity) | 0.15%<br>(2) | 0.74%<br>(2.68) | 0.01%<br>(2) | 0.01%<br>(2) | 20.49%<br>(2.34) |
|  | Jaccard | - | 0.91 | 0.94 | 0.94 | 0.92 |
|  | Fisher-test fdr | - | **<0.001** | 1 | 1 | **<0.001** |
| Gene symbol<br>↓<br>Ensembl GeneID | One to one | 78.44% | **81.78%** | 73.25% | 2.69% | 73.73% |
|  | Unmapped | **1.02%** | 7.98% | 7.4% | 97.22% | 6.35% |
|  | One to many<br>(Multiplicity) | 20.55%<br>(2.15) | 10.24%<br>(2.14) | 19.09%<br>(2.19) | 0.09%<br>(2) | 19.92%<br>(2.19) |
|  | Jaccard | - | 0.85 | 0.93 | 0.024 | 0.93 |
|  | Fisher test pval | - | **0.006** | **0.002** | **0.02** | **<0.001** |
| Entrez GeneID<br>↓<br>Gene symbol | One to one | **99.99%** | 88.46% | 96.97% | 96.92% | 67.72% |
|  | Unmapped | **0.01%** | 11.19% | 3.03% | 3.08% | 16.79% |
|  | One to many<br>(Multiplicity) | 0%<br>(-) | 0.36%<br>(2.51) | 0%<br>(-) | 0%<br>(-) | 15.49%<br>(2.16) |
|  | Jaccard | - | 0.85 | 0.97 | 0.98 | 0.82 |
|  | Fisher-test fdr | - | **<0.001** | 0.56 | 1 | **<0.001** |
| Entrez GeneID<br>↓<br>Ensembl GeneID | One to one | **77.55%** | 72.74% | 72.6% | 75.12% | 72.68% |
|  | Unmapped | **6.65%** | 10.42% | 10.28% | 10.63% | 10.57% |
|  | One to many<br>(Multiplicity) | 15.8%<br>(2.15) | 16.83%<br>(2.16) | 17.12%<br>(2.16) | 14.25%<br>(2.12) | 16.75%<br>(2.15) |
|  | Jaccard | - | 0.94 | 0.94 | 0.92 | 0.94 |
|  | Fisher-test fdr | - | **<0.001** | **<0.001** | **<0.001** | **<0.001** |
| Ensembl GeneID<br>↓<br>Gene symbol | One to one | **98.49%** | 93.95% | 87.39% | 95.38% | 74.15% |
|  | Unmapped | **0%** | 6.05% | 11.55% | 4.52% | 25.33% |
|  | One to many<br>(Multiplicity) | 1.51%<br>(2.04) | 0%<br>(-) | 1.06%<br>(3.76) | 0.1%<br>(4.52) | 0.51%<br>(2.23) |
|  | Jaccard | - | 0.87 | 0.86 | 0.91 | 0.74 |
|  | Fisher-test fdr | - | **<0.001** | **<0.001** | **<0.001** | **<0.001** |
| Ensembl GeneID<br>↓<br>Entrez GeneID | One to one | **89.53%** | 86.94% | 87.39% | 87.69% | 86.03% |
|  | Unmapped | **9.81%** | 11.93% | 11.55% | 12.21% | 13.2% |
|  | One to many<br>(Multiplicity) | 0.66%<br>(2.09) | 1.12%<br>(4.22) | 1.06%<br>(3.75) | 0.1%<br>(4.52) | 0.77%<br>(2.83) |
|  | Jaccard | - | 0.93 | 0.95 | 0.94 | 0.94 |
|  | Fisher-test fdr | - | **<0.001** | **<0.001** | **<0.001** | **<0.001** |

**Table S2.** Performance results of geneslator and the compared tools for gene identifiers mapping in *S*.*cerevisiae*.

| Mapping | *Saccharomyces cerevisiae* |  |  |  |  |  |
| --- | --- | --- | --- | --- | --- | --- |
|  |  | geneslator | biomaRt | Org.Sc.sgd.db | mygene | gprofiler2 |
| Gene symbol<br>↓<br>Entrez GeneID | One to one | **98.93%** | - | 0.32% | NA | 86.21% |
|  | Unmapped | **1.04%** | 100% | 99.68% | NA | 12.72% |
|  | One to many<br>(Multiplicity) | 0.03%<br>(2) | - | 0%<br>(-) | NA | 1.07%<br>(2.13) |
|  | Jaccard | - | 0 | 0.003 | - | 0.88 |
|  | Fisher-test fdr | - | - | 1 | - | 1 |
| Gene symbol<br>↓<br>Ensembl GeneID | One to one | 98.21% | - | - | NA | **100%** |
|  | Unmapped | 1.76% | 100% | 100% | NA | 0% |
|  | One to many<br>(Multiplicity) | 0.03%<br>(2) | - | - | NA | 0%<br>(-) |
|  | Jaccard | - | 0 | 0 | - | 0.98 |
|  | Fisher-test fdr | - | - | - | - | 1 |
| Entrez GeneID<br>↓<br>Gene symbol | One to one | **99.95%** | 85.12% | 90.04% | NA | 96.23% |
|  | Unmapped | **0.05%** | 13.44% | 9.96% | NA | 2.07% |
|  | One to many<br>(Multiplicity) | 0%<br>(-) | 1.45%<br>(2.13) | 0%<br>(-) | NA | 1.71%<br>(2.25) |
|  | Jaccard | - | 0.86 | 0.9 | - | 0.92 |
|  | Fisher-test fdr | - | 0.78 | 1 | - | **<0.001** |
| Entrez GeneID<br>↓<br>Ensembl GeneID | One to one | **99.9%** | 96.23% | 96.21% | NA | 96.23% |
|  | Unmapped | **0.07%** | 2.07% | 2.08% | NA | 2.07% |
|  | One to many<br>(Multiplicity) | 0.03%<br>(2) | 1.71%<br>(2.25) | 1.71%<br>(2.25) | NA | 1.71%<br>(2.25) |
|  | Jaccard | - | 0.98 | 0.98 | - | 0.98 |
|  | Fisher-test fdr | - | 1 | 1 | - | 1 |
| Ensembl GeneID<br>↓<br>Gene symbol | One to one | 96.97% | 76.3% | 77.08% | 0.03% | **99.96%** |
|  | Unmapped | **0.01%** | 23.7% | 21.68% | 99.97% | 0.04% |
|  | One to many<br>(Multiplicity) | 3.02%<br>(2) | 0%<br>(-) | 1.24%<br>(2.16) | 0%<br>(-) | 0%<br>(-) |
|  | Jaccard | - | 0.74 | 0.76 | 0 | 0.97 |
|  | Fisher-test fdr | - | 1 | 1 | **<0.001** | 1 |
| Ensembl GeneID<br>↓<br>Entrez GeneID | One to one | **96.88%** | 85.25% | 85.27% | 0.03% | 85.25% |
|  | Unmapped | **3.08%** | 13.3% | 13.33% | 99.97% | 13.3% |
|  | One to many<br>(Multiplicity) | 0.04%<br>(2) | 1.45%<br>(2.25) | 1.4%<br>(2.26) | 0%<br>(-) | 1.45%<br>(2.25) |
|  | Jaccard | - | 0.89 | 0.89 | 0 | 0.89 |
|  | Fisher-test fdr | - | 1 | 1 | **<0.001** | 1 |

**Table S3.** Performance results of geneslator and the compared tools for gene identifiers mapping in *A*.*thaliana*.

| Mapping | *Arabidopsis thaliana* |  |  |  |  |  |
| --- | --- | --- | --- | --- | --- | --- |
|  |  | geneslator | biomaRt | Org.At.tair.db | mygene | gprofiler2 |
| Gene symbol<br>↓<br>Entrez GeneID | One to one | **93.51%** | NA | 31.48% | 83.54% | 91.75% |
|  | Unmapped | **5.77%** | NA | 67.12% | 15.87% | 6.63% |
|  | One to many<br>(Multiplicity) | 0.72%<br>(2.06) | NA | 1.4%<br>(2.14) | 0.59%<br>(2.14) | 1.62%<br>(4.41) |
|  | JaccardI | - | NA | 0.35 | 0.89 | 0.90 |
|  | Fisher-test fdr | - | NA | **<0.001** | **<0.001** | **<0.001** |
| Gene symbol<br>↓<br>Ensembl GeneID | One to one | **93.68%** | NA | 32.14% | 45.91% | 92.67% |
|  | Unmapped | **5.43%** | NA | 66.41% | 53.77% | 5.72% |
|  | One to many<br>(Multiplicity) | 0.89%<br>(2.12) | NA | 1.44%<br>(2.15) | 0.32%<br>(2.09) | 1.61%<br>(2.33) |
|  | Jaccard | - | NA | 0.35 | 0.49 | 0.9 |
|  | Fisher-test fdr | - | NA | **<0.001** | **0.002** | **<0.001** |
| Entrez GeneID<br>↓<br>Gene symbol | One to one | **100%** | NA | 29.98% | 98.79% | 97.79% |
|  | Unmapped | **0%** | NA | 39.85% | 1.21% | 1.92% |
|  | One to many<br>(Multiplicity) | 0%<br>(-) | NA | 30.17%<br>(2.61) | 0%<br>(-) | 0.28%<br>(2.86) |
|  | Jaccard | - | NA | 0.26 | 0.99 | 0.39 |
|  | Fisher-test fdr | - | NA | **<0.001** | 1 | **<0.001** |
| Entrez GeneID<br>↓<br>Ensembl GeneID | One to one | **99.24%** | NA | 98.36% | NA | 97.83% |
|  | Unmapped | **0.47%** | NA | 1.64% | NA | 1.88% |
|  | One to many<br>(Multiplicity) | 0.29%<br>(2.17) | NA | 0%<br>(-) | NA | 0.29%<br>(2.92) |
|  | Jaccard | - | NA | 0.99 | NA | 0.99 |
|  | Fisher-test fdr | - | NA | 1 | NA | 1 |
| Ensembl GeneID<br>↓<br>Gene symbol | One to one | **99.72%** | NA | 23.4% | 81.74% | 83.94% |
|  | Unmapped | **0.03%** | NA | 55.27% | 18.22% | 15.79% |
|  | One to many<br>(Multiplicity) | 0.25%<br>(2) | NA | 21.33%<br>(2.59) | 0.04%<br>(2) | 0.27%<br>(3.01) |
|  | Jaccard | - | NA | 0.21 | 0.81 | 0.39 |
|  | Fisher-test fdr | - | NA | **<0.001** | 1 | **<0.001** |
| Ensembl GeneID<br>↓<br>Entrez GeneID | One to one | **99.14%** | NA | 79.57% | 84.17% | 80.98% |
|  | Unmapped | **0.60%** | NA | 20.43% | 15.79% | 18.67% |
|  | One to many<br>(Multiplicity) | 0.26%<br>(2.06) | NA | 0%<br>(-) | 0.04%<br>(2) | 0.35%<br>(10.64) |
|  | Jaccard | - | NA | 0.8 | 0.8 | 0.82 |
|  | Fisher-test fdr | - | NA | 1 | **<0.001** | 1 |

**Table S4.** Performance results of geneslator and the compared tools for gene identifiers mapping in *R*.*norvegicus*.

| Mapping |  | *Rattus norvegicus* |  |  |  |  |
| --- | --- | --- | --- | --- | --- | --- |
|  |  | geneslator | biomaRt | org.Rn.eg.db | mygene | gprofiler2 |
| Gene symbol<br>↓<br>Entrez GeneID | One to one | **99.43%** | 55.28% | 80.59% | 80.51% | 53.16% |
|  | Unmapped | **0.07%** | 44.01% | 19.41% | 19.49% | 44.43% |
|  | One to many<br>(Multiplicity) | 0.5%<br>(2.1) | 0.71%<br>(2.12) | 0%<br>(-) | 0%<br>(-) | 2.41%<br>(20.88) |
|  | Jaccard | - | 0.57 | 0.81 | 0.80 | 0.55 |
|  | Fisher-test fdr | - | **<0.001** | 1 | **0.02** | **<0.001** |
| Gene symbol<br>↓<br>Ensembl GeneID | One to one | **72.96%** | 56.66% | 60.83% | 28.53% | 59.99% |
|  | Unmapped | **25%** | 42.88% | 39.03% | 71.38% | 37.96% |
|  | One to many<br>(Multiplicity) | 2.04%<br>(2.15) | 0.46%<br>(2.02) | 0.14%<br>(9.57) | 0.09%<br>(2) | 2.05%<br>(2.48) |
|  | Jaccard | - | 0.77 | 0.80 | 0.38 | 0.73 |
|  | Fisher-test fdr | - | 0.21 | **<0.001** | **<0.001** | **<0.001** |
| Entrez GeneID<br>↓<br>Gene symbol | One to one | **99.98%** | 93.07% | 97.87% | 97.87% | 86.48% |
|  | Unmapped | **0.02%** | 6.89% | 2.13% | 2.13% | 10.84% |
|  | One to many<br>(Multiplicity) | 0%<br>(-) | 0.04%<br>(2.17) | 0%<br>(-) | 0%<br>(-) | 2.68%<br>(2.29) |
|  | Jaccard | - | 0.9 | 0.98 | 0.98 | 0.75 |
|  | Fisher-test fdr | - | **<0.001** | 1 | 1 | **<0.001** |
| Entrez GeneID<br>↓<br>Ensembl GeneID | One to one | **95.25%** | 93.6% | 93.99% | 39.28% | 89% |
|  | Unmapped | **3.91%** | 6.26% | 5.85% | 60.72% | 10.84% |
|  | One to many<br>(Multiplicity) | 0.84%<br>(2.03) | 0.14%<br>(2.16) | 0.16%<br>(2.13) | 0%<br>(-) | 0.16%<br>(2.09) |
|  | Jaccard | - | 0.968 | 0.972 | 0.408 | 0.882 |
|  | Fisher-test fdr | - | 0.632 | 0.627 | **<0.001** | **<0.001** |
| Ensembl GeneID<br>↓<br>Gene symbol | One to one | **97.76%** | 61.53% | 58.64% | 61.56% | 66.44% |
|  | Unmapped | **0.05%** | 38.47% | 40.9% | 38.42% | 34.39% |
|  | One to many<br>(Multiplicity) | 2.19%<br>(2.01) | 0%<br>(-) | 0.46%<br>(2.15) | 0.02%<br>(3) | 3.17%<br>(2.25) |
|  | Jaccard | - | 0.64 | 0.62 | 0.64 | 0.68 |
|  | Fisher-test fdr | - | 1 | **<0.001** | **<0.001** | **<0.001** |
| Ensembl GeneID<br>↓<br>Entrez GeneID | One to one | **66.78%** | 56.39% | 58.64% | 59.02% | 54.14% |
|  | Unmapped | **32.12%** | 43.18% | 40.9% | 40.96% | 45.19% |
|  | One to many<br>(Multiplicity) | 1.1%<br>(2.01) | 0.43%<br>(2.16) | 0.46%<br>(2.15) | 0.02%<br>(3) | 0.67%<br>(2.33) |
|  | Jaccard | - | 0.84 | 0.88 | 0.87 | 0.8 |
|  | Fisher-test fdr | - | **<0.001** | **<0.001** | **0.005** | **<0.001** |

**Table S5.** Performance results of geneslator and the compared tools for gene identifiers mapping in *D*.*melanogaster*.

| Mapping | *Drosophila melanogaster* |  |  |  |  |  |
| --- | --- | --- | --- | --- | --- | --- |
|  |  | geneslator | biomaRt | org.Dm.eg.db | mygene | gprofiler2 |
| Gene symbol<br>↓<br>Entrez GeneID | One to one | **98.77%** | 75.6% | 77.04% | 74.76% | 91.71% |
|  | Unmapped | **0.08%** | 22.97% | 22.96% | 25.24% | 6.88% |
|  | One to many<br>(Multiplicity) | 1.15%<br>(2.22) | 1.43%<br>(10.64) | 0%<br>(-) | 0%<br>(-) | 1.41%<br>(10.76) |
|  | Jaccard | - | 0.78 | 0.78 | 0.75 | 0.93 |
|  | Fisher-test fdr | - | 1 | 1 | 1 | 1 |
| Gene symbol<br>↓<br>Ensembl GeneID | One to one | **98.71%** | 76.81% | 75.77% | 71.97% | 92.96% |
|  | Unmapped | **0.08%** | 22.97% | 22.97% | 28.03% | 6.88% |
|  | One to many<br>(Multiplicity) | 1.21%<br>(2.23) | 0.22%<br>(2) | 1.26%<br>(11.8) | 0%<br>(-) | 0.16%<br>(2) |
|  | Jaccard | - | 0.78 | 0.78 | 0.72 | 0.93 |
|  | Fisher-test fdr | - | 1 | 0.55 | 1 | 1 |
| Entrez GeneID<br>↓<br>Ensembl GeneID | One to one | **99.82%** | 96.83% | 96.70% | 96.29% | 97.04% |
|  | Unmapped | **0%** | 1.84% | 1.97% | 3.71% | 1.63% |
|  | One to many<br>(Multiplicity) | 0.18%<br>(2) | 1.33%<br>(11.42) | 1.33%<br>(11.42) | 0%<br>(-) | 1.33%<br>(11.42) |
|  | Jaccard | - | 0.98 | 0.98 | 0.96 | 0.98 |
|  | Fisher-test fdr | - | 1 | 1 | 1 | 1 |
| Entrez GeneID<br>↓<br>Gene symbol | One to one | **100%** | 96.83% | 98.03% | 98.02% | 97.03% |
|  | Unmapped | **0%** | 1.84% | 1.97% | 1.98% | 1.64% |
|  | One to many<br>(Multiplicity) | 0%<br>(-) | 1.33%<br>(11.42) | 0%<br>(-) | 0%<br>(-) | 1.33%<br>(11.42) |
|  | Jaccard | - | 0.98 | 0.98 | 0.98 | 0.95 |
|  | Fisher-test fdr | - | 1 | 1 | 1 | **0.002** |
| Ensembl GeneID<br>↓<br>Gene symbol | One to one | **99.86%** | 99.28% | 96.52% | 98.89% | 99.5% |
|  | Unmapped | **0.13%** | 0.72% | 2.32% | 1.11% | 0.5% |
|  | One to many<br>(Multiplicity) | 0.01%<br>(2) | 0%<br>(-) | 1.16%<br>(11.32) | 0%<br>(-) | 0%<br>(-) |
|  | Jaccard | - | 0.99 | 0.98 | 0.99 | 0.97 |
|  | Fisher-test fdr | 1 | 1 | 1 | 1 | **0.22** |
| Ensembl GeneID<br>↓<br>Entrez GeneID | One to one | **99.74%** | 96.91% | 96.52% | 98.88% | 97.13% |
|  | Unmapped | **0.25%** | 1.93% | 2.32% | 1.12% | 1.71% |
|  | One to many<br>(Multiplicity) | 0.01%<br>(2) | 1.16%<br>(11.32) | 1.16%<br>(11.32) | 0%<br>(-) | 1.16%<br>(11.32) |
|  | Jaccard | - | 0.97 | 0.98 | 0.99 | 0.98 |
|  | Fisher-test fdr | - | 1 | 1 | 1 | 1 |

**Table S6.** Performance results of geneslator and the compared tools for gene identifiers mapping in *C*.*elegans*.

| Mapping | *Caenorhabditis elegans* |  |  |  |  |  |
| --- | --- | --- | --- | --- | --- | --- |
|  |  | geneslator | biomaRt | org.Ce.eg.db | mygene | gprofiler2 |
| Gene symbol<br>↓<br>Entrez GeneID | One to one | **99.61%** | 75.88% | 78.36% | 77.74% | 89.86% |
|  | Unmapped | **0.38%** | 23.7% | 21.64% | 22.26% | 5.23% |
|  | One to many<br>(Multiplicity) | 0.01%<br>(2) | 0.42%<br>(2.11) | 0%<br>(-) | 0%<br>(-) | 4.91%<br>(2.91) |
|  | Jaccard | - | 0.77 | 0.79 | 0.78 | 0.91 |
|  | Fisher-test fdr | - | 0.45 | 1 | 1 | **<0.001** |
| Gene symbol<br>↓<br>Ensembl GeneID | One to one | **98.73%** | 75.91% | 74.51% | 74.79% | 92.50% |
|  | Unmapped | **0.79%** | 23.68% | 24.65% | 25.2% | 2.01% |
|  | One to many<br>(Multiplicity) | 0.48%<br>(2.12) | 0.41%<br>(2.1) | 0.84%<br>(2.19) | 0.01%<br>(2) | 5.49%<br>(2.25) |
|  | Jaccard | - | 0.77 | 0.76 | 0.75 | 0.94 |
|  | Fisher-test fdr | - | 1 | 1 | 1 | **<0.001** |
| Entrez GeneID<br>↓<br>Gene symbol | One to one | **100%** | 92.33% | 98.44% | 98.44% | 55.21% |
|  | Unmapped | **0%** | 7.42% | 1.56% | 1.56% | 44.46% |
|  | One to many<br>(Multiplicity) | 0%<br>(-) | 0.25%<br>(2.6) | 0%<br>(-) | 0%<br>(-) | 0.33%<br>(3.45) |
|  | Jaccard | - | 0.88 | 0.98 | 0.98 | 0.55 |
|  | Fisher-test fdr | - | **<0.001** | 0.24 | 1 | **<0.001** |
| Entrez GeneID<br>↓<br>Ensembl GeneID | One to one | **98.66%** | 89.64% | 89.64% | 92.67% | 89.64% |
|  | Unmapped | **0.02%** | 7.42% | 7.42% | 7.33% | 7.42% |
|  | One to many<br>(Multiplicity) | 1.32%<br>(2.11) | 2.94%<br>(2.54) | 2.94%<br>(2.54) | 0%<br>(-) | 2.94%<br>(2.54) |
|  | Jaccard | - | 0.93 | 0.93 | 0.93 | 0.93 |
|  | Fisher-test fdr | - | 1 | 1 | 1 | 1 |
| Ensembl GeneID<br>↓<br>Gene symbol | One to one | **97.09%** | 93.84% | 90.83% | 93.94% | 57.43% |
|  | Unmapped | **0%** | 6.16% | 6.17% | 6.06% | 42% |
|  | One to many<br>(Multiplicity) | 2.91%<br>(2) | 0%<br>(-) | 3%<br>(2.53) | 0%<br>(-) | 0.57%<br>(2.35) |
|  | Jaccard | - | 0.91 | 0.92 | 0.93 | 0.57 |
|  | Fisher-test fdr | - | 1 | 1 | 1 | **<0.001** |
| Ensembl GeneID<br>↓<br>Entrez GeneID | One to one | **98.42%** | 90.83% | 90.83% | 93.91% | 89.42% |
|  | Unmapped | **0.1%** | 6.17% | 6.17% | 6.09% | 6.07% |
|  | One to many<br>(Multiplicity) | 1.48%<br>(2) | 3%<br>(2.53) | 3%<br>(2.53) | 0%<br>(-) | 4.51%<br>(2.47) |
|  | Jaccard | - | 0.94 | 0.94 | 0.94 | 0.94 |
|  | Fisher-test fdr | - | 1 | 1 | 1 | 1 |

## Footnotes

1 https://bioconductor.org/packages/release/data/annotation/

## References

1. Yunze Liu and Gang Li. Empowering biologists to decode omics data: the genekitr r package and web server. BMC Bioinformatics, 24(1):1–14, 2023.

2. Michael Ashburner, Catherine A. Ball, Judith A. Blake, David Botstein, Heather Butler, J. Michael Cherry, Allan P. Davis, Kara Dolinski, Selina S. Dwight, Janan T. Eppig, et al. Gene ontology: tool for the unification of biology. Nature Genetics, 25(1):25–29, 2000.

3. Minoru Kanehisa, Mika Furumichi, Mao Tanabe, Yoko Sato, and Kanae Morishima. Kegg: new perspectives on genomes, pathways, diseases and drugs. Nucleic Acids Research, 45(D1):D353–D361, 2017.

4. Imre Vastrik, Peter D’Eustachio, Esther Schmidt, Geeta Joshi-Tope, Gopal Gopinath, David Croft, Bernard de Bono, Marc Gillespie, Bijay Jassal, Suzanna Lewis, et al. Reactome: a knowledge base of biologic pathways and processes. Genome Biology, 8(3):R39, 2007.

5. Anjali Agrawal et al. Wikipathways 2024: next generation pathway database. Nucleic Acids Research, 52(D1):D679–D687, 2024.

6. G. R. Brown, V. Hem, K. S. Katz, et al. Gene: a gene-centered information resource at ncbi. Nucleic Acids Research, 43(D1):D36–D42, 2015.

7. Ruth L Seal, Bryony Braschi, Kristian Gray, Tamsin EM Jones, Susan Tweedie, Liora Haim-Vilmovsky, and Elspeth A Bruford. Genenames. org: the hgnc resources in 2023. Nucleic Acids Research, 51(D1):D1003–D1009, 2023.

8. Peter W Harrison, M Ridwan Amode, Olanrewaju Austine-Orimoloye, Andrey G Azov, Matthieu Barba, If Barnes, Arne Becker, Ruth Bennett, Andrew Berry, Jyothish Bhai, et al. Ensembl 2024. Nucleic acids research, 52(D1):D891–D899, 2024.

9. Shifu Chen and Chunlei Wu. mygene: R client for MyGene.info services, 2016. R package version 1.24.0.

10. Liis Kolberg, Uku Raudvere, Ivan Kuzmin, Jaak Vilo, and Hedi Peterson. gprofiler2– an r package for gene list functional enrichment analysis and namespace conversion toolset g:profiler. F1000Research, 9 (ELIXIR)(709), 2020. R package version 0.2.4.

11. The UniProt Consortium. Uniprot: a hub for protein information. Nucleic Acids Research, 43(Database issue):D204–D212, 2015.

12. J. T. Eppig et al. Mouse genome informatics (mgi): knowledgebase for laboratory mouse. ILAR Journal, 58(1):17–41, 2017.

13. Mahima Vedi, Jennifer R Smith, G Thomas Hayman, Monika Tutaj, Kent C Brodie, Jeffrey L De Pons, Wendy M Demos, Adam C Gibson, Mary L Kaldunski, Logan Lamers, Stanley J F Laulederkind, Jyothi Thota, Ketaki Thorat, Marek A Tutaj, Shur-Jen Wang, Stacy Zacher, Melinda R Dwinell, and Anne E Kwitek. 2022 updates to the rat genome database: a findable, accessible, interoperable, and reusable (fair) resource. Genetics, 224(1):iyad042, May 2023.

14. S. R. Engel, S. Aleksander, R. S. Nash, E. D. Wong, S. Weng, S. R. Miyasato, G. Sherlock, and J. M. Cherry. Saccharomyces genome database: Advances in genome annotation, expanded biochemical pathways, and other key enhancements. Genetics, page iyae185, November 2024.

15. Paul W. Sternberg, Kimberly Van Auken, Qinghua Wang, Adam Wright, Karen Yook, Magdalena Zarowiecki, Valerio Arnaboldi, and Andrés Becerra. Wormbase 2024: status and transitioning to alliance infrastructure. Genetics, 227(1):iyae050, May 2024.

16. Arzu Öztürk Çolak, Steven J Marygold, Giulia Antonazzo, Helen Attrill, Damien Goutte-Gattat, Victoria K Jenkins, Beverley B Matthews, Gillian Millburn, Gilberto dos Santos, Christopher J Tabone, and FlyBase Consortium. Flybase: updates to the drosophila genes and genomes database. Genetics, 227(1):iyad211, May 2024.

17. Y. M. Bradford, C. E. Van Slyke, L. Ruzicka, A. Singer, A. Eagle, D. Fashena, D. G. Howe, K. Frazer, R. Martin, H. Paddock, C. Pich, S. Ramachandran, and M. Westerfield. Zebrafish information network, the knowledgebase for danio rerio research. Genetics, 220(4), 2022.

18. Leonore Reiser, Erica Bakker, Sabarinath Subramaniam, Xingguo Chen, Swapnil Sawant, Kartik Khosa, Trilok Prithvi, and Tanya Z. Berardini. The arabidopsis information resource in 2024. Genetics, 227(1):iyae027, May 2024.

19. Tina A. Eyre, Mathew W. Wright, Michael J. Lush, and Elspeth A. Bruford. Hcop: a searchable database of human orthology predictions. Briefings in Bioinformatics, 8(1):2–5, January 2007.

20. Alliance of Genome Resources Consortium. Updates to the alliance of genome resources central infrastructure. Genetics, 227(1):iyae049, May 2024.

21. Adrian M. Altenhoff et al. OMA orthology in 2024: improved prokaryote coverage, ancestral and extant GO enrichment, a revamped synteny viewer and more in the OMA ecosystem. Nucleic Acids Research, 52(D1):D513–D521, 2024.

22. Salvatore Cosentino and Wataru Iwasaki. SonicParanoid: fast, accurate and easy orthology inference. Bioinformatics, 35(1):149–151, 2019.

23. Erik L.L. Sonnhammer and Mateusz Kaduk. InParanoiDB 9: Ortholog groups for protein domains and full-length proteins. Journal of Molecular Biology, 435(14):168001, 2023.

24. Fergal J. Martin et al. Ensembl 2023. Nucleic Acids Research, 51(D1):D933–D941, 2023.

25. Yannis Nevers, Arnaud Kress, Audrey Defosset, Raymond Ripp, Benjamin Linard, Julie D. Thompson, Olivier Poch, and Odile Lecompte. OrthoInspector 3.0: open portal for comparative genomics. Nucleic Acids Research, 47(D1):D411–D418, 2019.

26. Diego Fuentes, Manuel Molina, Uciel Chorostecki, Salvador Capella-Gutiérrez, Marina Marcet-Houben, and Toni Gabaldón. PhylomeDB v5: an expanding repository for genome-wide catalogues of annotated gene phylogenies. Nucleic Acids Research, 50(D1):D1062–D1068, 2022.

27. Paul D. Thomas, Dustin Ebert, Anushya Muruganujan, Tremayne Mushayahama, Laurent-Philippe Albou, and Huaiyu Mi. PANTHER: Making genome-scale phylogenetics accessible to all. Protein Science, 31(1):8–22, 2022.

28. David M. Emms and Steven Kelly. OrthoFinder: phylogenetic orthology inference for comparative genomics. Genome Biology, 20(1):238, 2019.

29. Mateusz Kaduk, Christian Riegler, Oliver Lemp, and Erik L.L. Sonnhammer. HieranoiDB: a database of orthologs inferred by Hieranoid. Nucleic Acids Research, 45(D1):D687–D690, 2017.

30. Ayushi Agrawal, Hasan Balcai, and Kristina et al. Hanspers. Wikipathways 2024: next generation pathway database. Nucleic Acids Research, 52(D1):D679–D689, 11 2023.

31. M. Carlson and H. Pagès. AnnotationForge: Tools for building SQLite-based annotation data packages, 2025. R package version 1.52.0, available from: https://bioconductor.org/packages/AnnotationForge.

32. Hervé Pagès, Marc Carlson, Seth Falcon, and Nianhua Li. AnnotationDbi: Manipulation of SQLite-based annotations in Bioconductor, 2025. R package version 1.73.0.

33. L. C. Sommerfeld, J. Schrapers, K. F. Müller, L. Bravo-Merodio, et al. High rate triggers increased atrial release of bmp10, a biomarker for atrial fibrillation and stroke, and bmp10 affects ventricular cardiomyocytes. Circulation: Arrhythmia and Electrophysiology, 18(11):e013834, November 2025.

34. Antonio Colaprico, Tiago C Silva, Catharina Olsen, Luciano Garofano, Claudia Cava, Davide Garolini, Thais S Sabedot, Tathiane M Malta, Stefano M Pagnotta, Isabella Castiglioni, et al. Tcgabiolinks: an r/bioconductor package for integrative analysis of tcga data. Nucleic acids research, 44(8):e71–e71, 2016.

35. Aparna R Parikh, Annamaria Szabolcs, Jill N Allen, Jeffrey W Clark, Jennifer Y Wo, Michael Raabe, Hannah Thel, David Hoyos, Arnav Mehta, Sanya Arshad, et al. Radiation therapy enhances immunotherapy response in microsatellite stable colorectal and pancreatic adenocarcinoma in a phase ii trial. Nature cancer, 2(11):1124–1135, 2021.

36. F. Wang, X. Liu, S. Li, C. Zhao, et al. Resolving the lineage relationship between malignant cells and vascular cells in glioblastomas. Protein & Cell, 14(2):105–122, March 2023.

37. A. Olander, C. M. Ramirez, V. H. Acosta, P. Medina, et al. Pregnancy reduces il33+ hybrid progenitor accumulation in the aged mammary gland. bioRxiv, August 2024. Preprint.

38. Z. Ren, P. Yu, D. Li, Z. Li, et al. Single-cell reconstruction of progression trajectory reveals intervention principles in pathological cardiac hypertrophy. Circulation, 141(21):1704–1719, May 2020.

39. P. Liu, H. Zhang, C. A. Werley, M. Pichler, S. Ryan, C. Lewarch, J. Jacques, J. Grooms, J. Ferrante, G. Li, D. Zhang, N. Bremmer, A. Barnett, R. Chantre, A. E. Elder, A. E. Cohen, L. A. Williams, G. T. Dempsey, and O. B. McManus. A phenotypic screening platform for chronic pain therapeutics using all-optical electrophysiology. Gene Expression Omnibus (GEO), August 2023. GEO accession: GSE237797.

40. E. Cid, A. Marquez-Galera, M. Valero, B. Gal, et al. Sublayer- and cell-type-specific neurodegenerative transcriptional trajectories in hippocampal sclerosis. Cell Reports, 35(10):109229, June 2021.

41. S. Lee, J. N. Chun, H. J. Lee, H. H. Park, et al. Transcriptome analysis of the anti-tgf*β* effect of Schisandra chinensis fruit extract and schisandrin b in a7r5 vascular smooth muscle cells. Life, 11(2), February 2021.

42. D. H. Kim, D. Grün, and A. van Oudenaarden. Dampening of expression oscillations by synchronous regulation of a microrna and its target. Nature Genetics, 45(11):1337–1344, November 2013.

43. S. T. Page and R. Haluch. Whole animal mrna profiles of *C. elegans* nematodes with and without *bir-2* knockdown and *E. faecalis*. Gene Expression Omnibus (GEO), September 2021. Unpublished dataset; submission date Sep 14, 2021; last updated Dec 31, 2022.

44. E. S. Yang, S. Y. Cheon, J. Y. Park, Y. Park, et al. Ilimaquinone-induced lipophagy diminishes lipid accumulation via ampk activation. BMB Reports, 58(9):415–423, September 2025.

45. K. Narbonne-Reveau, E. Lanet, C. Dillard, S. Foppolo, et al. Neural stem cell-encoded temporal patterning delineates an early window of malignant susceptibility in drosophila. eLife, 5, June 2016.

46. N. A. Tchurikov, E. S. Klushevskaya, D. M. Fedoseeva, I. R. Alembekov, et al. Dynamics of whole-genome contacts of nucleoli in drosophila cells suggests a role for rdna genes in global epigenetic regulation. Cells, 9(12), December 2020.

47. X. Xu, J. Gao, G. Shan, L. Zhang, et al. Dibromoacetonitrile exposure induces neurobehavioral toxicity in adult zebrafish via disruption of multiple neural pathways. Biochemical and Biophysical Research Communications, 786:152732, October 2025.

48. R. Dorsky, D. Vasudevan, and S. Alper. Unpublished dataset from neurobiology lab: [details not specified], August 2024. Submission date: Aug 14, 2024; last update: Aug 15, 2025.

49. P. Moura-Alves, A. Puyskens, A. Stinn, M. Klemm, et al. Host monitoring of quorum sensing during pseudomonas aeruginosa infection. Science, 366(6472), December 2019.

50. T. Turgut Genç and M. Günay. Saccharomyces cerevisiae by4741 raw sequence reads. NCBI BioProject PRJNA1177667, October 2024. Submission date: Oct 28, 2024; last update: Oct 01, 2025.

51. I. Iermak, N. R. Wilson Eisele, K. Kurscheidt, M. J. Loukeri, et al. Molecular mechanisms governing the formation of distinct upf1-containing complexes in yeast. Cell Reports, 44(10):116415, October 2025.

52. Affymetrix, Inc. Affymetrix arabidopsis genechip annotation and support resources. http://www.affymetrix.com/support/technical/byproduct.affx?product=arab, July 2002. Submission date: Jul 01, 2002; last update: Jun 12, 2017. Annotation updates: NetAffx build 28 (2009), build 32 (2012), build 35 (2016). Contact.

53. T. Barros-Galvão, X. Chen, and S. Penfield. Unpublished dataset from crop genetics lab: [details not specified]. NCBI Gene Expression Omnibus (GEO), March 2025. Submission date: Mar 19, 2025; last update: Oct 08, 2025.

54. C. Bergis-Ser, Q. Wang, X. He, M. Nisa, V. Kaiser, C. Mazubert, J. Drouin-Wahbi, R. Brik-Chaouche, L. Chmaiss, J. Van Leene, G. de Jaeger, J. Gutierrez-Marcos, C. Bergounioux, C. Richet-Bourbousse, D. Latrasse, M. Benhamed, and C. Raynaud. Rna-seq datasets integrating spike-in and classic experiments in arabidopsis thaliana mutants. NCBI Gene Expression Omnibus (GEO), March 2025. Submission date: Mar 25, 2025; last update: Nov 18, 2025.

55. Steffen Durinck, Paul T. Spellman, Ewan Birney, and Wolfgang Huber. biomaRt: Interface to BioMart databases (i.e. Ensembl), 2005. R package version 2.54.0.

56. Marc Carlson. org.Hs.eg.db: Genome wide annotation for Human, 2025. R package version 3.22.0.

57. Marc Carlson. org.Mm.eg.db: Genome wide annotation for Mouse, 2025. R package version 3.22.0.

58. Marc Carlson. org.Rn.eg.db: Genome wide annotation for Rat, 2025. R package version 3.22.0.

59. Marc Carlson. org.Dm.eg.db: Genome wide annotation for Fly, 2025. R package version 3.22.0.

60. Marc Carlson. org.Ce.eg.db: Genome wide annotation for Worm, 2025. R package version 3.22.0.

61. Marc Carlson. org.Dn.eg.db: Genome wide annotation for Zebrafish, 2025. R package version 3.22.0.

62. Marc Carlson. org.Sc.sgd.db: Genome wide annotation for Yeast, 2025. R package version 3.22.0.

63. Marc Carlson. org.At.tair.db: Genome wide annotation for Arabidopsis, 2025. R package version 3.22.0.

64. Brian M. Schilder and Nathan G. Skene. orthogene: Interspecies Gene Mapping, 2022. R package version 1.4.0.

65. Ogan Mancarci and Leon French. homologene: Quick Access to Homologene and Gene Annotation Updates, 2023. R package version 1.7.

66. Igor Dolgalev. babelgene: Gene Orthologs for Model Organisms in a Tidy Data Format, 2022. R package version 22.9.

67. Liis Kolberg, Uku Raudvere, et al. gprofiler2 – an r package for gene list functional enrichment analysis and identifier conversion. PLoS ONE, 15(3), 2020.

68. Guangchuang Yu, Liguang Wang, Yanyan Han, and Qing-Yu He. clusterprofiler: an r package for comparing biological themes among gene clusters. OMICS: A Journal of Integrative Biology, 16(5):284–287, 2012.

69. Elena Pontarini et al. Serum and tissue biomarkers associated with composite of relevant endpoints for Sjögren syndrome (CRESS) and Sjögren tool for assessing response (STAR) to B cell-targeted therapy in the trial of anti-B cell therapy in patients with primary Sjögren syndrome (TRACTISS). Arthritis & Rheumatology, 76(5):763–776, May 2024.

70. Matthew E Ritchie, Belinda Phipson, D. Wu, Yifang Hu, Charity W Law, Wei Shi, and Gordon K Smyth. limma powers differential expression analyses for RNA-sequencing and microarray studies. Nucleic Acids Research, 43(7):e47, 2015.

71. Yunshun Chen, Lizhong Chen, Aaron T L Lun, Pedro Baldoni, and Gordon K Smyth. edgeR v4: powerful differential analysis of sequencing data with expanded functionality and improved support for small counts and larger datasets. Nucleic Acids Research, 53(2):gkaf018, 2025.

